# Development of spike receptor-binding domain nanoparticle as a vaccine candidate against SARS-CoV-2 infection in ferrets

**DOI:** 10.1101/2021.01.28.428743

**Authors:** Young-Il Kim, Dokyun Kim, Kwang-Min Yu, Hogyu David Seo, Shin-Ae Lee, Mark Anthony B. Casel, Seung-Gyu Jang, Stephanie Kim, WooRam Jung, Chih-Jen Lai, Young Ki Choi, Jae U. Jung

## Abstract

Severe acute respiratory syndrome coronavirus 2 (SARS-CoV-2), a causative agent of COVID-19 pandemic, enters host cells *via* the interaction of its Receptor-Binding Domain (RBD) of Spike protein with host Angiotensin-Converting Enzyme 2 (ACE2). Therefore, RBD is a promising vaccine target to induce protective immunity against SARS-CoV-2 infection. In this study, we report the development of RBD protein-based vaccine candidate against SARS-CoV-2 using self-assembling *H. pylori*-bullfrog ferritin nanoparticles as an antigen delivery. RBD-ferritin protein purified from mammalian cells efficiently assembled into 24-mer nanoparticles. 16-20 months-old ferrets were vaccinated with RBD-ferritin nanoparticles (RBD-nanoparticles) by intramuscular or intranasal inoculation. All vaccinated ferrets with RBD-nanoparticles produced potent neutralizing antibodies against SARS-CoV-2. Strikingly, vaccinated ferrets demonstrated efficient protection from SARS-CoV-2 challenge, showing no fever, body weight loss and clinical symptoms. Furthermore, vaccinated ferrets showed rapid clearance of infectious viruses in nasal washes and lungs as well as viral RNA in respiratory organs. This study demonstrates the Spike RBD-nanoparticle as an effective protein vaccine candidate against SARS-CoV-2.

## Introduction

SARS-CoV-2, originally named 2019-nCoV upon initial isolation from Wuhan, China in December 2019, has caused a global outbreak of coronavirus disease-19 (COVID-19) with significant socioeconomic impacts (1, 2). From the continuously growing numbers of diagnoses and deaths, COVID-19 was declared a public health emergency of international concern (PHEIC) in January 2020 and soon declared a pandemic by WHO in March 2020 (3, 4). As of Jan 27^th^ 2021, more than 100 million people have been infected with SARS-CoV-2, among which 2 million died (5). Although approximately 80% of the confirmed SARS-CoV-2 infections are asymptomatic or show mild flu-like symptoms, 20% of the infections progress to severe pneumonia and acute respiratory distress syndrome requiring hospitalization and mechanical ventilation (6, 7). The overwhelming number of SARS-CoV-2 patients has rapidly devastated the availability of health-care resources (8). Shortage of medical resources and staff in conjunction with the overwhelming number of patients have exacerbated the quality of medical care and eventually increased mortality rates of COVID-19 (9). Although a significant proportion of the infected patients have recovered, many of them report cardiovascular, pulmonary, and neurologic symptoms lasting after the recovery (10, 11). Thus, strong preventive measures are essential to halt the pandemic and its destructive effects on the global public health, as well as the economy.

SARS-CoV-2 is a member of the *Coronaviridae* family, carrying a single positive-stranded RNA genome within the viral envelope (2). Although at least seven coronaviruses are known as etiological agents of mild respiratory illnesses in human infection, the family has not been closely associated with severe illnesses until the relatively recent outbreaks of SARS-CoV, MERS-CoV, and SARS-CoV-2 (1, 12). Emergence of these pathogens and the COVID-19 pandemic have called for an urgent global research efforts to investigate the pathogenesis of coronaviruses. The SARS-CoV-2 RNA genome is approximately 30 kilobases and encodes for structural proteins such as–Spike (S), Envelope (E), Membrane (M), and Nucleocapsid (N)– and non-structural proteins such as papain-like protease, chymotrypsin-like protease, and RNA-dependent RNA polymerase (13). The heavily glycosylated S protein protruding from the virion surface is the key bridge between the virus and the host cell, playing a crucial role in host cell receptor recognition, virion attachment, and ultimately entry into the host cell. S is a member of the Class I viral fusion protein which undergoes trimerization upon cleavage into S1 and S2 domains by a host cellular protease, furin. While S1 confers specificity in cell tropism through its Receptor-Binding Domain (RBD) which directly interacts with the receptor of SARS-CoV-2, Angiotensin-Converting Enzyme 2 (ACE2), S2 mediates membrane fusion via formation of a trimeric hairpin structure from its heptad repeat domains (14). Therefore, S1 RBD has been considered as one of the most promising candidates in vaccine development to protect against Coronaviruses (15–17). Its efficacy has previously been shown to induce potent neutralizing antibodies against MERS-CoV (18). Furthermore, previous studies of neutralizing antibodies from naturally recovered patients of SARS-CoV-2 infections have mapped their epitopes to be S1 and RBD (19, 20), implicating RBD-targeting antibodies in successful immunity against SARS-CoV-2 (21–23). Thus, most of the currently developed vaccines against SARS-CoV-2,–despite of their diversity in vaccine approaches,–include RBD in their immunogens (24–28).

One major limitation of small soluble proteins alone as vaccine candidates is that our immune system only reacts efficiently against immunogens of nanometer range in size (29, 30). Therefore, many protein vaccines using viral proteins are developed into virus-like particles (VLPs) which are multiprotein structures that mimic the organization and conformation of native viruses, but lack the viral genome. However, this approach is limited to a few pathogens that are capable of self-assembling into VLPs upon overexpression of the viral protein, such as Hepatitis B virus (HBV) surface antigen (HBsAg) and Human Papillomavirus (HPV) L1 protein (31–33). Fortunately, the latest advances in molecular biology and nanotechnology have overcome this limitation by adopting nanoparticle engineering to serve as platforms for vaccines. The efficacy of these nanoparticle-engineered vaccines outperforms traditional vaccines, such as whole inactivated vaccines of bacterial and viral pathogens (34–38). Moreover, recent studies have shed light on the immunological advantages of nanoparticle-based vaccines in nearly every step of the humoral and cellular immunity: efficient antigen transport to draining lymph nodes, antigen presentation by follicular dendritic and helper T cells, as well as high level of activation of the germinal centers (30, 39, 40). Among the genetically engineered nanoparticles, ferritin is the most well-characterized in the bionanotechnology field. Ferritin, ubiquitous through kingdoms of life, has a conserved role in minimizing damage to cell from reactive oxygen species formed from the Fenton reaction upon excess iron (II). Due to its natural tendency to self-assemble into 24-meric homopolymer and amenability via fusion peptides, ferritin is an ideal candidate for drug delivery and vaccine development (41, 42). Most importantly, its exceptional chemical and thermal stability does not require stringent temperature control, enabling streamlined distribution process, especially in areas with limited resources for cold-chain supplies (41, 43). One of the recently engineered ferritin for vaccine development is the self-assembling *Helicobacter pylori*-bullfrog (*Rana Catesbeiana*) hybrid ferritin which carries NH_2_-terminal residues from the lower subunit of bullfrog ferritin on the core of *Helicobacter pylori* ferritin to form radially projecting tails (38). The *H. pylori* ferritin-based nanoparticle has been reported to be an effective platform for vaccines to carry trimeric glycoproteins for presenting viral immunogens on its threefold axis points. Most importantly, it provides stronger protective immunity at a lower dose than soluble immunogens against Influenza and Epstein-Barr viruses, while minimizing the risk of autoimmunity through its genetic diversity from heavy and light chains of human ferritin (38, 44, 45).

Despite recent efforts to develop mouse models that fully recapitulates human SARS-CoV-2 infection, the current hACE2-transgenic mouse model fails to mimic pathogenic progress and symptoms of COVID-19 in humans. Ferrets (*Mustela putorius furo*) on the other hand, are naturally susceptible to human respiratory viruses–Respiratory Syncytial virus (48), Influenza virus (49, 50), and SARS-CoV (51, 52)–making ferret models ideal to study respiratory virus infections in humans. In addition, ferrets share anatomy of upper and lower respiratory tracts, architecture of terminal bronchioles, and density of submucosal glands to those of humans (46, 47). Recently, we and others have shown that SARS-CoV-2-infected ferrets develop immune responses and pathogenic progress similar to humans’, and shed virus through nasal wash, saliva, urine, and fecal samples, which highly recapitulate human SARS-CoV-2 infection (53–56). Furthermore, we have also demonstrated the efficacy of the ferret model in drug discovery for SARS-CoV-2 (57). Thus, ferrets represent an infection and transmission animal model of SARS-CoV-2 that should facilitate the development of SARS-CoV-2 therapeutics and vaccines.

Here, we demonstrate the immunogenic efficacy of self-assembling spike RBD-ferritin nanoparticle (RBD-nanoparticle) as an efficient SARS-CoV-2 vaccine antigen. We purified the RBD-nanoparticle from transfected HEK293T cells and immunized ferrets via intramuscular (IM) and intranasal (IN) routes to monitor the induction of neutralizing antibodies. Furthermore, we challenged the vaccinated ferrets with SARS-CoV-2 and observed protective immunity against SARS-CoV-2. Taken together, we propose the self-assembling RBD-nanoparticles as a potential vaccine candidate that effectively protects against SARS-CoV-2 infection.

## Results

### Purification and characterization of RBD-ferritin nanoparticles

Kanekiyo et al. have discovered the use of engineered ferritin in vaccine developments by fusion with viral immunogens (38, 44). Briefly, the NH_2_-terminal tail from the lower subunit of bullfrog ferritin was fused to *H. pylori* ferritin so that the bullfrog-originated tail and viral immunogen were fused by the linker and presented on the threefold axis points of the *H. pylori* ferritin core. The human codon-optimized RBD of SARS-CoV-2 Wuhan-Hu-1 strain (NC_045512) was fused to the IL-2 signal peptide at the amino terminus and the *H. pylori*-bullfrog ferritin at the carboxyl terminus to generate the RBD-ferritin fusion. A computer-assisted modeling predicts the 3D structure of RBD-ferritin nanoparticles with RBD forming radial projections on the threefold axis point of fully assembled nanoparticles (Fig. 1A). Ferritin and RBD-ferritin fusion proteins were readily purified from the supernatants of transfected HEK293T cells (Fig. 1B). To demonstrate the 24-mer self-assembly of ferritin nanoparticles, purified ferritin and RBD-ferritin proteins were subjected to size exclusion chromatography with columns designed to have a maximum resolution for proteins with kilodalton and megadalton ranges of molecular weight. As a result, the purified ferritin nanoparticles and RBD-nanoparticles showed peaks at approximately 408 kDa and 1350 kDa, respectively, corresponding to 24-mers of each protein (Fig. 1C). These results indicate that RBD-ferritin protein is readily purified from mammalian cells to homogeneity and efficiently assembles into 24-mer nanoparticles.

**Fig 1.**
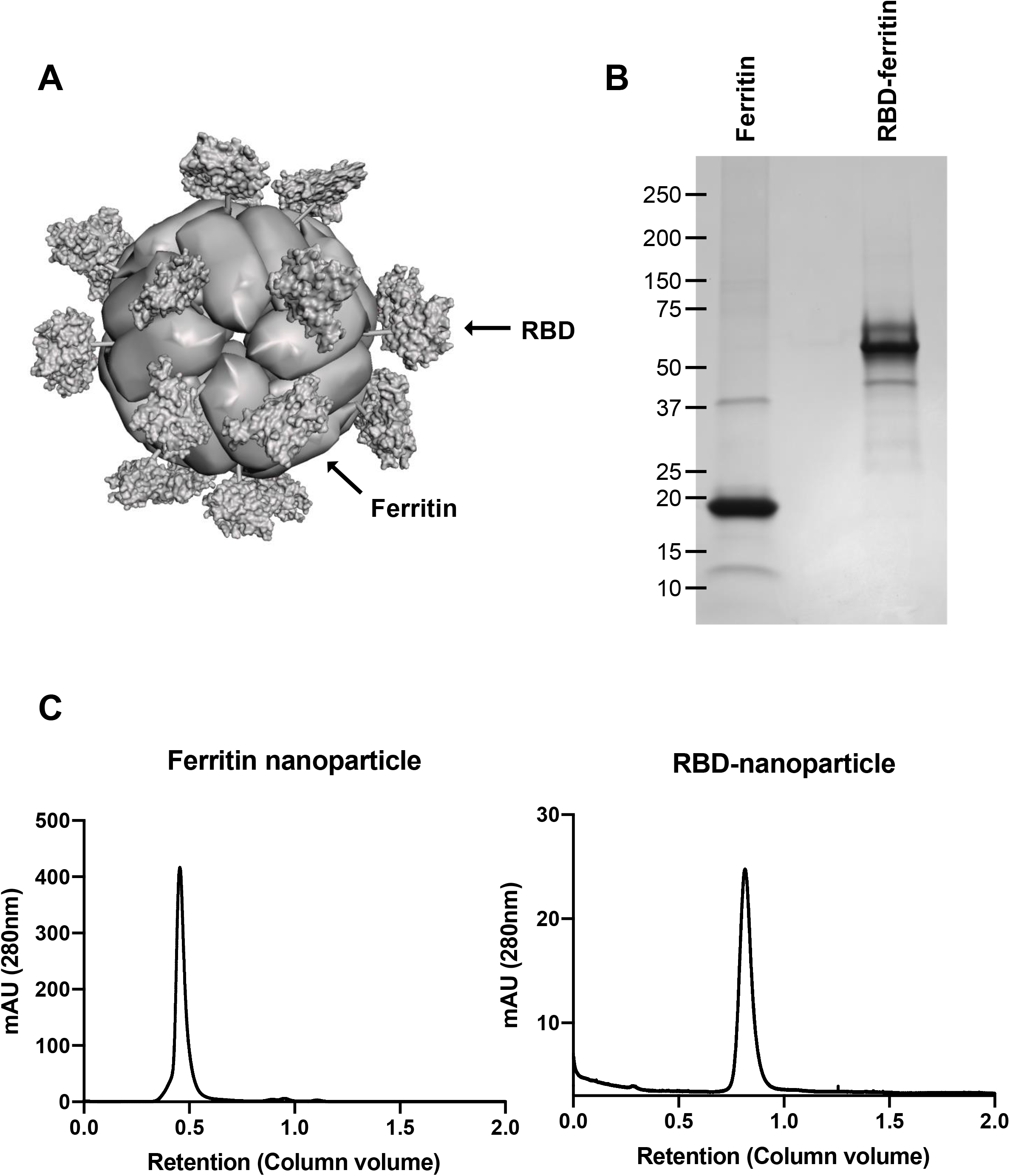
Design and purification of RBD-nanoparticle. A. Computer-assisted modeling of RBD-nanoparticle based on previously solved structures of *H. pylori* ferritin (PDB: 3EGM) and SARS-CoV-2 RBD (PDB: 7JMP). RBD forms radial projections on threefold axis point of fully assembled nanoparticle. B. Coomassie staining of purified ferritin-nanoparticle and RBD-nanoparticle following SDS-PAGE. C. Size exclusion chromatography peaks of the concentrated supernatants from HEK293T transfected with plasmids encoding secreted ferritin-nanoparticle or RBD-nanoparticles. The supernatants were concentrated with 100 kDa MWCO and 500 kDa MWCO filters on TFF system and loaded to Superdex 200 Increase 10/300 GL and HiPrep 16/60 Sephacryl S-500 HR gel filtration columns on Bio-rad NGC chromatography system, respecitvely.

### Immunization with RBD-nanoparticle induces neutralizing antibody in ferrets

To test the vaccine efficacy of purified RBD-nanoparticles, we immunized 16-20 months-old ferrets (n=10/immunization route), which is equivalent to 30 years of age in humans. While intramuscular (IM) immunization is the most widely used route for vaccine delivery, intranasal (IN) immunization closely resembles infection with respiratory pathogens and efficiently stimulates mucosal immunity (58). Ferrets were injected with 15μg RBD-nanoparticle via IM route only or both IM and IN routes over 31 days with boosting immunizations at days 14 and 28 (Fig. 2A). Blood was drawn from each ferret prior to primary and boosting immunizations on days 14 and 28. All ferrets vaccinated with RBD-nanoparticles produced strong neutralizing antibodies after the second boosting immunization performed at day 28. Neutralization titer did not show statistically significant difference between the routes of immunization (Fig. 2B). These data indicate that RBD-nanoparticle immunization induces strong neutralizing antibody regardless of the route of immunization.

**Fig 2.**
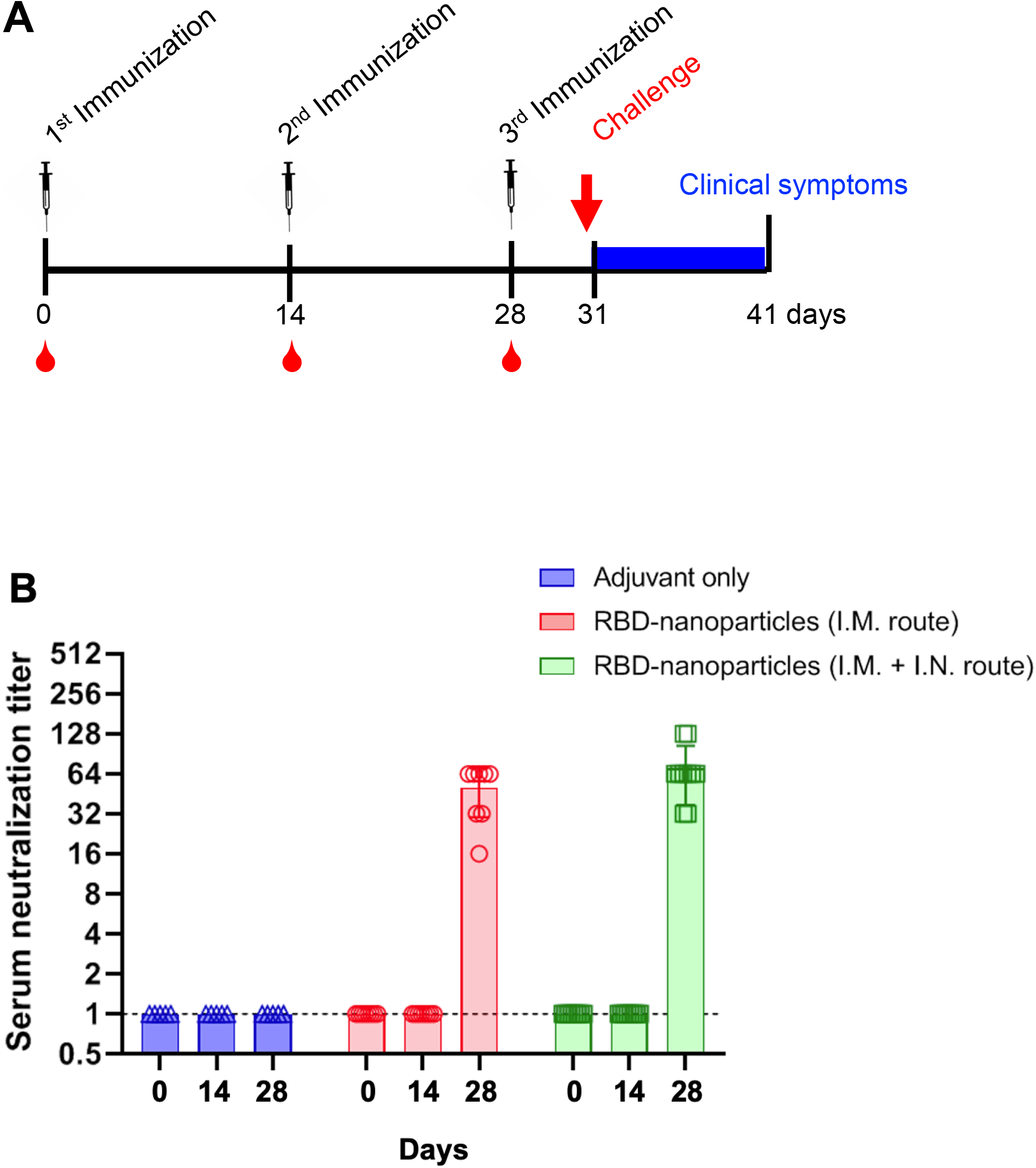
Immunization with RBD-nanoparticle elicits neutralizing antibody formation. A. Immunization schedule of ferrets. At day 31, ferrets were challenged with 10^5.0^ TCID_50_/mL of SARS-CoV-2 and observed for clinical symptoms for the following 10 days. One group was immunized with only PBS and adjuvant (only adjuvant-immunized), and two other groups were immunized with 15μg RBD-nanoparticle in adjuvant with 1:1 ratio for total volume of 600μl. B. Serum neutralization titer of adjuvant-immunized, RBD-nanoparticle IM immunized, or RBD-nanoparticle IM and IN immunized ferrets. Neutralizing antibody titers against SARS-CoV-2 NMC2019-nCoV02 (100 TCID_50_) of ferritin-nanoparticle immunized groups were measured in Vero cells with serially diluted ferret sera collected before immunizations at days 0, 14 and 28.

### Immunization with RBD-nanoparticle promotes rapid viral clearance and protects ferrets from SARS-CoV-2 challenge

Immunized ferrets were challenged with 10^5.0^ TCID_50_/mL of NMC2019-nCoV02 strain SARS-CoV-2 three days after the last immunization at day 31 and monitored for clinical symptoms resembling COVID-19. Ferrets with adjuvant only-immunization was included as control group. Over a total of 10 days from the day of challenge infection, ferrets with adjuvant only-immunization showed increase in body temperature and decrease in body weight (Fig. 3A). On the contrary, ferrets immunized with RBD-nanoparticle did not show any change in either body temperature or body weight (Fig. 3A and B). Minor body weight changes in ferrets immunized by IM route showed a statistically insignificant difference compared to the adjuvant only-immunized ferrets (Fig. 3B). On the other hand, ferrets immunized by IM and IN routes provided stronger protection with high statistical significance against body weight loss as shown by the minimal reduction of body weight followed by a constant increase thereafter (Fig. 3B). Nasal wash samples were collected every other day for 10 days after the virus challenge, and 3 ferrets were sacrificed at 3 and 6 days post-infection (dpi) to harvest the lungs. Consistent with the trend shown in body temperature and weight, immunized ferrets showed rapid viral clearance in nasal washes (Fig. 3C) and lungs (Fig. 3D) of both groups of vaccinated ferrets. It should be noted that IM and IN immunization showed slightly more effective viral clearance in nasal washes at 4 dpi than IM immunization (Fig. 3C).

**Fig 3.**
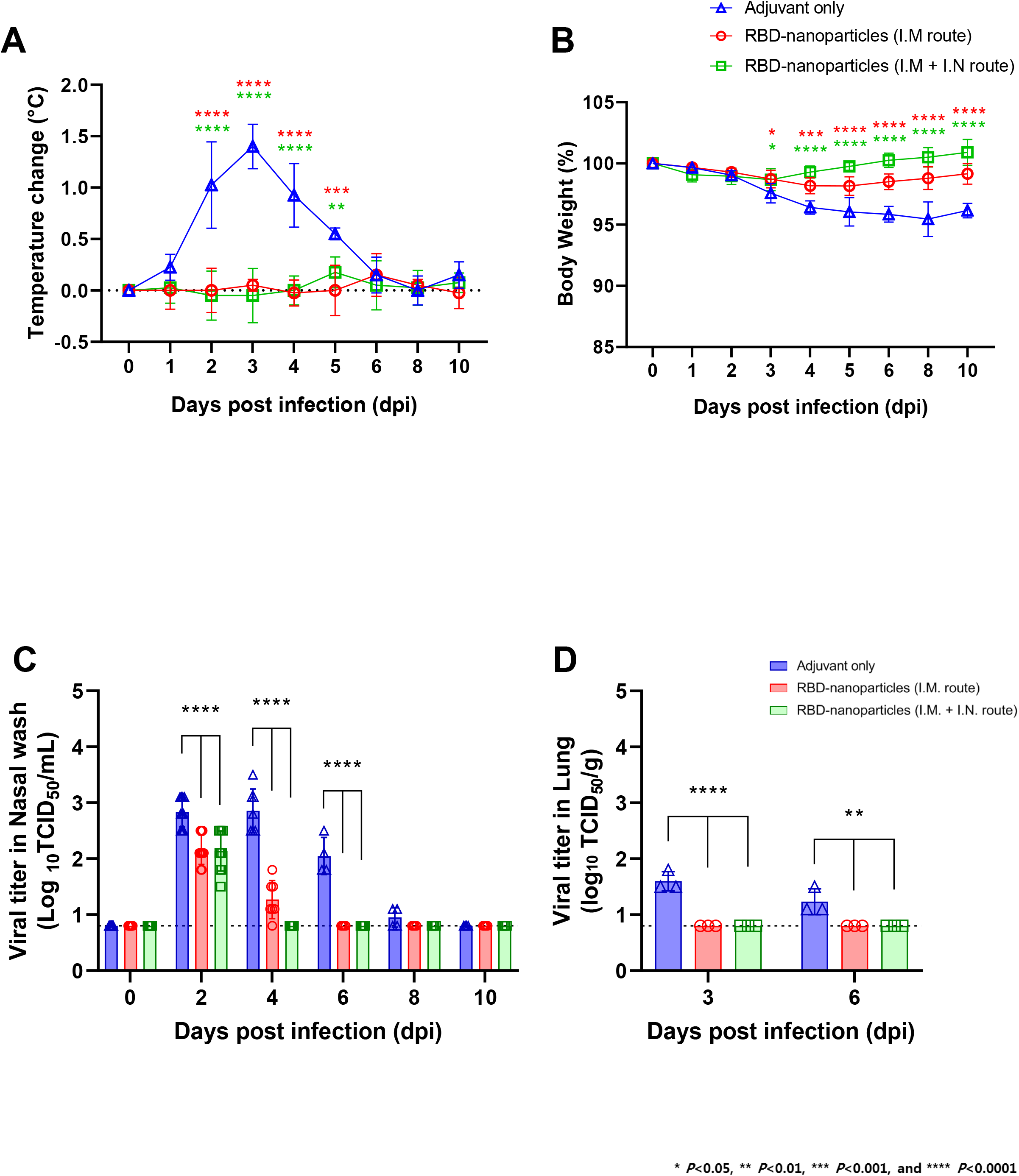
Immunization with RBD-nanoparticle promotes rapid viral clearance and protects ferrets from SARS-CoV-2 challenge. A. Body temperature change of adjuvant-immunized, RBD-nanoparticle IM immunized, or RBD-nanoparticle IM and IN immunized ferrets upon SARS-CoV-2 challenge. B. Body weight change of adjuvant-immunized, RBD-nanoparticle IM immunized, or RBD-nanoparticle IM and IN immunized ferrets upon SARS-CoV-2 challenge. C. Viral titer in the nasal washes of adjuvant-immunized, RBD-nanoparticle IM immunized, or RBD-nanoparticle IM and IN immunized ferrets upon SARS-CoV-2 challenge. D. Viral titer in the lung tissue homogenates of adjuvant-immunized, RBD-nanoparticle IM immunized, or RBD-nanoparticle IM and IN immunized ferrets upon SARS-CoV-2 challenge.

To further investigate the potency of protective immunity by RBD-nanoparticle, we challenged the immunized ferrets with a higher titer (10^6.0^ TCID_50_/mL) of SARS-CoV-2 following the same immunization protocol (Fig. 2A). Consistently, RBD-nanoparticles-immunized ferrets showed no increase of body temperature compared to adjuvant only-immunized ferrets (Fig. S1). While adjuvant only-immunized ferrets suffered from cough, runny nose, and reduction in movement, RBD-nanoparticles-immunized ferrets showed only mild reduction in movement on the 2^nd^ and 3^rd^ days after the high virus titer challenge (Table 1). On the other hand, IN and IM immunization showed more potent protective immunity upon challenge with a high virus titer than IN immunization only (Fig. S2). IN and IM immunization led to faster clearance of infectious virus in nasal washes at 4 and 8 dpi than IM immunization alone (Fig. S2A). Infectious virus titers of lungs were also lower in IN and IM-immunized ferrets than in IM-immunized ferrets (Fig. S2B). These data demonstrate that RBD-nanoparticle induces strong protective immunity to suppress SARS-CoV-2-induced clinical symptoms and promote viral clearance. Moreover, a combination of IN and IM immunization induces stronger anti-viral immunity against challenge of high titer SARS-CoV-2 than IM immunization alone.

**Table 1.** RBD-nanoparticle immunization suppresses clinical symptoms induced by challenge with high SARS-CoV-2 titer.

| Group<br>(n = 4/group) | Clinical | 0 dpi | 1 dpi | 2 dpi | 3 dpi | 4 dpi | 5 dpi | 6 dpi | 7 dpi | 8 dpi | 10 dpi |
| --- | --- | --- | --- | --- | --- | --- | --- | --- | --- | --- | --- |
| **Adjuvant**<br>**only** | Cough | 0 | 0 | 0.5 | 1 | 1 | 0 | 0 | 0 | 0 | 0 |
|  | Runny nose | 0 | 0 | 1.0 | 1 | 1 | 1 | 1 | 0.75 | 0 | 0 |
|  | Movement, activity | 0 | 0 | 1.25 | 2 | 2 | 1.25 | 0.75 | 0.5 | 0 | 0 |
|  | Total | 0 | 0 | 2.75 | 4 | 4 | 2.25 | 1.75 | 1.25 | 0 | 0 |
| **RBD-**<br>**nanoparticle** | Cough | 0 | 0 | 0 | 0 | 0 | 0 | 0 | 0 | 0 | 0 |
|  | Runny nose | 0 | 0 | 0 | 0 | 0 | 0 | 0 | 0 | 0 | 0 |
|  | Movement, activity | 0 | 0 | 0.75 | 0.5 | 0 | 0 | 0 | 0 | 0 | 0 |
|  | Total | 0 | 0 | 0.75 | 0.5 | 0 | 0 | 0 | 0 | 0 | 0 |

**A group of adjuvant-immunized or RBD-nanoparticle IM immunized ferrets were challenged with $10^{6.0}$ TCID<sub>50</sub>/mL of SARS-CoV-2 and observed for their clinical symptoms – cough, runny nose, movement, and activity. The symptoms were quantified as counts per 30 minutes.**

### RBD-ferritin vaccination blocks lung damage from SARS-CoV-2 challenge

COVID-19 has most commonly been shown to be associated with a spectrum of lung damage. To compare lung histopathologies among immunized ferrets, RNAscope *in situ* hybridization and histopathological examination were conducted (Fig. 4). Lung tissues harvested from naïve ferrets were included as negative controls (Fig. 4D). RNAscope *in situ* hybridization results showed that the adjuvant only-immunized ferrets had a number of SARS-CoV-2 RNA-positive cells at 3 and 6 dpi with infiltration of numerous inflammatory immune cells (Fig. 4A-E). At 3 dpi, IM- or IM and IN-immunized ferrets showed considerable reduction of viral RNAs in the lungs compared to adjuvant only-immunized ferrets (Fig. 4). At 6 dpi, lung tissues of IM- or IM and IN-immunized ferrets showed complete clearance of viral RNAs (Fig. 4F-G), while adjuvant only-immunized ferrets still showed high viral RNAs (Fig. 4E). Finally, IM- or IM and IN-immunized ferrets showed little or no infiltration of inflammatory immune cells in infected lung (Fig. 4B-G). These data show that RBD-nanoparticle immunization accelerates viral clearance in the lung and suppresses the infiltration of inflammatory immune cells.

**Fig 4.**
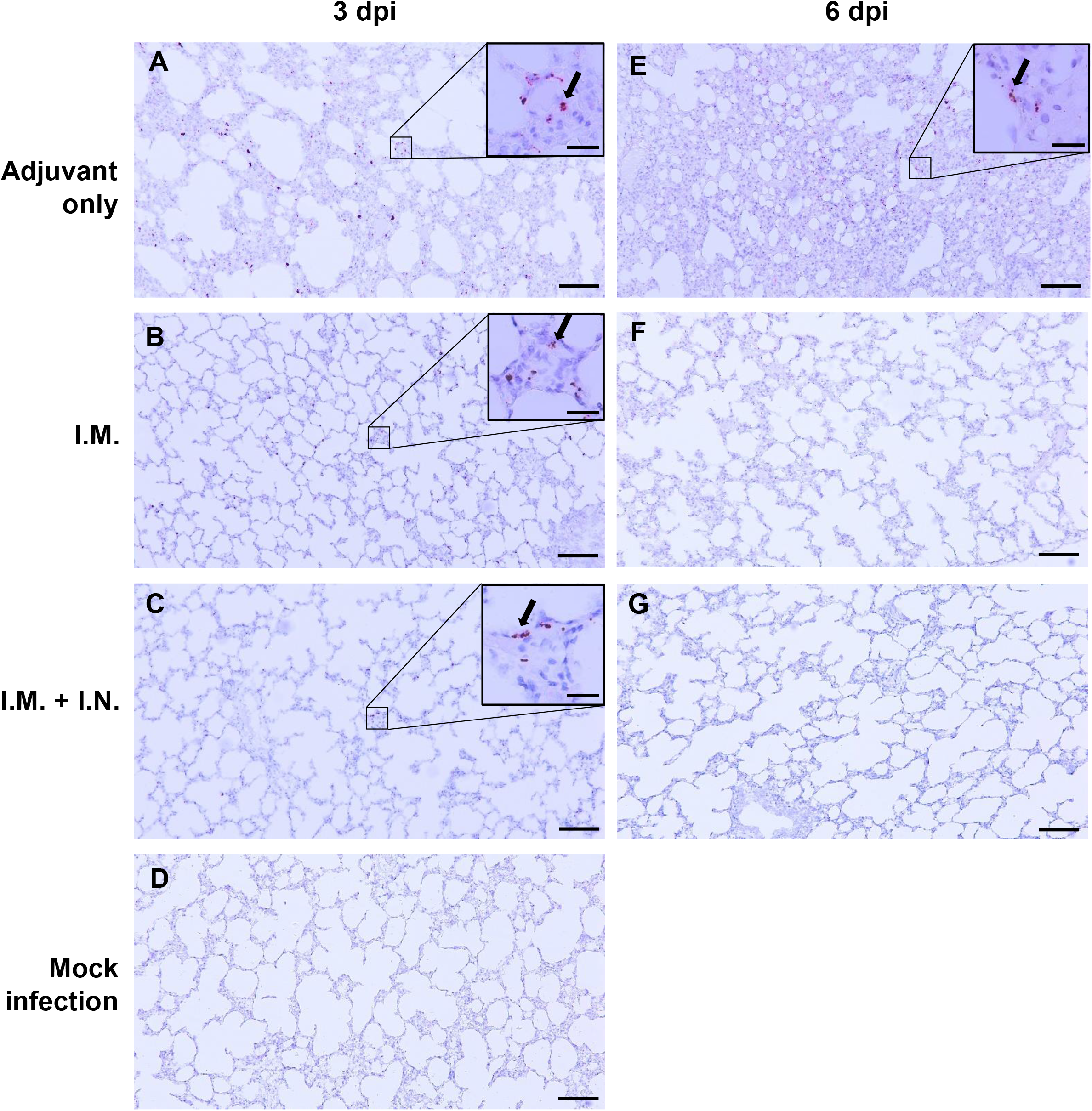
Lung histology and RNAscope of immunized ferrets upon SARS-CoV-2 challenge. Adjuvant-immunized, RBD-nanoparticle IM immunized, or RBD-nanoparticle IM and IN immunized ferrets were intranasally inoculated with 10^5.0^ TCID_50_/mL of SARS-CoV-2. Tissues were harvested on 3 and 6 dpi. RNAscope detected SARS-CoV-2 Spike RNA-positive cells in lung tissues of adjuvant-immunized (A and E), RBD-nanoparticle IM immunizated (B and F), and RBD-nanoparticle IM and IN immunizated ferrets (C and G). Mock infected ferret lung (D) was included as control. Magnification is x100 and scale bars represents 100 μm. Insert indicates the magnification (x400) of SARS-CoV-2-positive image and scale bar represents 20 μm. Black arrow indicates SARS-CoV-2 RNA-positive cells.

## Discussion

Since the first discovery in Wuhan, China in late 2019, SARS-CoV-2 has rapidly spread around the world and was declared a pandemic in 3 months. Confirmed infection and death counts have skyrocketed to over 88 million infections and 2 million deaths, and the statistics are still on a continuous rise. Although 80% of the infections do not progress to severe COVID-19, the recent surge in infections and severe patients have led to subsequent increase in mortality rates (8, 9). While several vaccines were approved at accelerated rates (59, 60), additional indepth study of mRNA-based vaccines regarding safety concerns and long-term effects still need to be addressed as they are the first approved mRNA human vaccine of its kind. Moreover, taking the growing evidence of reinfections into consideration, recovered patients cannot be completely excluded from the population requiring vaccination (61–64). Therefore, there still is a constant need for alternative vaccine approaches against SARS-CoV-2 using relatively well-characterized approaches. Recent advances in nanotechnology has favorably allowed the application of nanoparticles in the field of vaccinology to develop safer yet potent vaccines. One of the most promising candidates is *H. pylori*-bullfrog ferritin that has been genetically engineered to carry a protruding tail from the bullfrog on the self-assembling ferritin core of *H. pylori*, and serves as a platform to build nanoparticles of immunogen. This approach has proven higher efficacy in lower dose than traditional protein subunit vaccines. This approach also highlights lower risk of vaccine-related adverse effects and potentially greater accessibility to the public with reduced production cost (38, 44, 45). Importantly, inherent stability of ferritin nanoparticles from heat and chemicals may shed light to remove the necessity of strict cold-chain supply required for the currently distributing mRNA-based vaccines (41).

SARS-CoV-2 carries Spike protein to attach to the host receptor ACE2, which triggers membrane fusion for entry into host cells. The RBD of Spike protein confers the specificity to bind ACE2 and therefore is a promising target for vaccine development throughout the *Coronaviridae* family. As also shown from previously developed vaccine candidates against Coronaviruses (17, 65, 66), we selected the RBD as a vaccine antigen. However, soluble antigen is weakly immunogenic and therefore require higher dose of antigen along with an adjuvant, which correlates with higher risk of vaccine-related adverse effects (29). In this study, we engineered the fusion of SARS-CoV-2 Spike RBD with *H. pylori*-bullfrog ferritin to develop RBD-nanoparticle vaccine. Ferrets immunized with RBD-nanoparticles carried efficient neutralizing antibodies against SARS-CoV-2 and were protected from fever and body weight loss upon SARS-CoV-2 challenge. These clinical symptoms corresponded to the accelerated viral clearance in nasal washes and lungs following SARS-CoV-2 challenge. We further investigated the vaccine potential of RBD-nanoparticles by challenging the immunized ferrets with a high virus titer (10^6.0^ TCID_50_/mL). Immunized ferrets showed considerably reduced clinical symptoms, such as body weight loss, cough, runny nose, and movement activity, upon challenge of high titer SARS-CoV-2. Moreover, RNAscope analyses showed rapid viral clearance in the lungs of immunized ferrets compared to those of adjuvant only-immunized ferrets. Histological analysis also showed little or no lung tissue damage and inflammatory immune cell infiltration in immunized ferrets. As seen from other protein vaccines such as HPV VLP that requires prime-boost regimens (67), the first immunization alone was not sufficient to induce neutralizing antibodies. IN + IM immunization elicited more potent protective immunity upon challenge of high titer SARS-CoV-2 than IM immunization alone, which is consistent with previous reports showing stronger induction of mucosal immunity upon IN than IM immunization to protect against respiratory pathogens such as MERS-CoV (65, 66, 68), Influenza virus (69), and *Mycoplasma pneumoniae* (70). To differentiate vaccine efficacy between IN immunization and IN and IM immunization, we repeated the viral challenge with high titer (10^6.0^ TCID_50_/mL) and observed the improvement of viral clearance in lung and nasal washes from IN and IM-immunized ferrets. However, as IN and IM immunization was employed together in this study, further investigation is required to directly compare vaccine efficacy between IN immunization and IM immunization against SARS-CoV-2 infection. Also, ferrets challenged with 10^6.0^ TCID_50_/mL virus titer showed delayed viral clearance compared to ferrets challenged with 10^5.0^ TCID_50_/mL virus titer. However, 10^5.0^ TCID_50_/mL is already excessive and not physiologically relevant to real clinical setting.

In this study, we integrated SARS-CoV-2-derived immunogen into self-assembling nanoparticle to develop an effective vaccine candidate against COVID-19. IM-immunized animals showed strong induction of neutralizing antibody, rapid clearance of respiratory track virus, and clear suppression of clinical symptoms, which is further enhanced in combination with intranasal immunization. However, additional comprehensive studies are needed to understand the humoral and cellular immunity elicited by RBD-nanoparticle administration and differential activation of IgA-mediated mucosal immunity upon different immunization routes. Taken together, our study indicated that immunization with self-assembling SARS-CoV-2 RBD-nanoparticle elicits protective immunity against SARS-CoV-2 infection, showing its potential as a vaccine candidate in the midst of the COVID-19 pandemic.

## Material and Methods

### Material and Reagents

| REAGENT or RESOURCE | VENDOR | CATALOG NO. |
| --- | --- | --- |
| **Recombinant DNA** |  |  |
| pFUSEN-hIgG1Fc | Invivogen | pfcn-hg1 |
| *H. pylori*-bullfrog recombinant ferritin | Jefferey Cohen and Gary Nabel at NIAID |  |
| SARS-CoV-2 Spike codon-optimized to human codon usage | Genscript | MC_0101081 |
| **Chemicals** |  |  |
| Polyethylene imine | Polysciences | 23966 |
| Valproic acid | Sigma | P4543 |
| AddaVax adjuvant | Invivogen | Vac-adx-10 |
| RNAscope reagent | ACD | 322360 |
| RNAscope probe | ACD | 848561 |
| Gill's Hematoxylin #1 | Polysciences | 24242 |
| **Purification** |  |  |
| Labscale TFF system | Sigma | C1975 |
| TFF filter – 100 kDa MWCO | Sigma | PXB100C50 |
| TFF filter – 500 kDa MWCO | Sigma | PXB500C50 |
| NGC Medium-pressure liquid chromatography system | Bio-rad |  |
| BioFracFraction collector | Bio-rad | 7410002 |
| Superdex 200 Increase 10/300 GL | Cytiva | 45-002-570 |
| HiPrep 16/60 Sephacryl S-500 HR | Cytiva | 28935606 |

### Expression vector construction

The gene encoding the recombinant ferritin engineered from *Helicobacter pylori* nonheme ferritin and 2^nd^ to 9^th^ residues of bullfrog (*Rana catesbeiana*) ferritin lower subunit was a gift from Gary Nabel (44). Gene encoding Spike of SARS-CoV-2 (GenBank NC_0101080) codon-optimized for human codon usage (GenBank MC_0101081) was purchased from Genscript (pUC57-2019-nCoV-S). RBD was used to generate a fragment encoding RBD-SSGGASVLA linker-recombinant ferritin. For expression plasmid, a commercially available pFUSE vector (Invivogen) was engineered to replace human ferritin light chain gene promoter with SV40 promoter. Genes encoding the recombinant ferritin and the RBD-linker-ferritin fragment were cloned into the plasmid vector.

### Computer-assisted three-dimensional model of nanoparticles

Previously solved structures of *H. pylori* ferritin nanoparticle (PDB: 3EGM) and SARS-CoV-2 RBD (PDB: 7JMP) were processed with PyMol (Schrodinger) and Autodesk Meshmixer (Autodesk). The model was generated to reflect the linker connecting the end of RBD to the start of *H. pylori* ferritin monomer.

### Expression and purification of nanoparticles

HEK293T cells were directly purchased from American Type Culture Collection (ATCC) and maintained in DMEM medium (Gibco) supplemented with 10% FBS (Gibco) and 1% penicillin/streptomycin (Gibco). The cells were transiently transfected with polyethylenimine (Polysciences) and respective vector plasmids in Opti-MEM and FreeStyle 293 medium (Gibco) supplemented with 3mM valproic acid. Supernatants containing the nanoparticle were harvested 72 h after transfection and concentrated with Labscale TFF system equipped with 100 kDa and 500 kDa MWCO filters (Millipore Sigma). The concentrates were purified by size exclusion chromatography (NGC Medium-Pressure Liquid Chromatography, Bio-Rad) using Superdex 200 10/300 GL and HiPrep 16/60 Sephacryl S-500 HR (Cytiva) running degased PBS at 0.4ml/min. Standard curves were plotted using Gel filtration LMW/HMW calibration Kit (Cytiva) running at same conditions. Collected fractions were verified for their yield and purity via SDS-PAGE and stored at −80°C in 10% glycerol (Invitrogen).

### Virus propagation

NMC2019-nCoV02 strain of SARS-CoV-2 was isolated from a patient diagnosed with COVID-19 and tested positive for SARS-CoV-2 in February, 2020 in South Korea. Vero cells were used to propagate the virus in DMEM medium (Gibco) supplemented with 1% penicillin/streptomycin (Gibco) at 37°C. The viruses were harvested 72 h later and stored at − 80°C until use.

### Animal Care

Male and female ferrets of 16-20 months old and tested seronegative for Influenza A, MERS-CoV, and SARS-CoV were purchased from ID Bio Corporation (Cheongju, Korea). The ferrets were housed in ABSL3 facility within Chungbuk National University (Cheongju, Korea) with 12 h light/dark cycle with access to water and diet. All animal cares were performed strictly following the animal care guideline and experiment protocols approved by the Institutional Animal Care and Use Committee (IACUC) in Chungbuk National University.

### Ferret immunizations and viral challenge

RBD-ferritin nanoparticles (volume: 300ul) and AddaVax adjuvant (volume: 300ul) were administered into the legs through intramuscular injection and/or intranasal route. Subsequently, ferrets were intranasally infected with 10^5.0^ or 10^6.0^ TCID_50_/mL SARS-CoV-2. Body weight and temperature were measured, and veterinary clinical symptoms were observed every day. Blood and nasal washes were collected every other day for 10 days. Three animals per group were sacrificed at days 3 and 6 to collect lung tissues with individual scissors. Infectious viruses from the nasal washes and lung tissues were quantified by inoculation onto Vero cells. Veterinary symptoms were scored accordingly to our previous publication (57).

### Titration of neutralizing antibody in serum

The neutralizing antibody assay against SARS-CoV-2 was carried out using a microneutralization assay in Vero cells. Collected ferret serum specimens were inactivated at 56°C for 30 min. Initial 1:2 serum dilutions were made with the medium, and two-fold serial dilutions of all samples were made to a final serum dilution of 1:2 to 1:256. For each well, 50 μL of serially diluted serum was mixed with 50 μL (equal volume) of 100 TCID_50_ of SARS-CoV-2 and incubated at 37°C for 1 h to neutralize the infectious virus. The mixtures were then transferred to Vero cell monolayers. Vero cells were incubated at 37°C in 5% CO2 for 4 days and monitored for 50% reduction in cytopathic effect (CPE).

### RNAscope

SARS-CoV-2 RNA (Spike gene) was detected using the Spike-specific probe (Advanced Cell Diagnostics, Cat. # 848561) and visualized using RNAscope 2.5 HD Reagent Kit RED (Advanced Cell Diagnostics, Cat. # 322360). Lung tissue sections were fixed in 4% neutral-buffered formalin and embedded in paraffin, according to the manufacturer’s instructions, followed by counterstaining with 50% Gill’s hematoxylin #1 (Polysciences, cat # 24242-1000). Slides were viewed using Olympus IX 71 (Olympus, Tokyo, Japan) microscope with DP controller software to capture images.

### Statistical Analysis

All figure asterisks indicate statistical significance compared with adjuvant-only group as evaluated by the two-way ANOVA Dunnett’s multiple comparisons tests (* indicates p<0.05, ** indicates p<0.01, *** indicates p<0.001 and **** indicates p<0.0001) and were drawn using GraphPad Prism 8 (GraphPad).

## Acknowledgments

*H. pylori*-bullfrog ferritin construct was kindly provided by Drs. Jeffrey Cohen and Gary Nabel at Vaccine Research Center, NIAID. This work was supported by the National Research Foundation of Korea (NRF-2020R1A5A2017476, 2020R1A2C3008339), Korea Research Institute of Bioscience and Biotechnology (KRIBB) Research Initiative Program (KGM9942011), and National Institute of Health (CA200422, CA251275, AI140705, AI140705S, AI140718, AI152190, AI116585, AI116585S, DE023926, DE027888, and DE028521).

## Author Contributions

Y.I.K, D.K., Y.K.C. and J.U.J. conceived the study and designed the experiments. Y.I.K. and D.K. performed the experiments. K.M.Y., H.S., S.A.L., M.A.C., S.G.J., S.K., W.J. and C.J.L. helped with the experimental designs and data interpretation/analysis. Y.I.K. and D.K. took the lead to prepare the manuscript with Y.K.C. and J.U.J.

## Competing Financial Interests

Dr. Jae U Jung is a scientific adviser of the Vaccine Stabilization Institute, a California corporation.

---

**Fig S1. Body temperature of RBD-nanoparticle immunized ferrets against challenge with high titer SARS-CoV-2**

**Body temperature change of adjuvant-immunized, RBD-nanoparticle IM immunized, or RBD-nanoparticle IM and IN immunized ferrets upon high titer SARS-CoV-2 challenge.**

**Fig S2. Respiratory virus titer of RBD-nanoparticle immunized ferrets against challenge with high titer SARS-CoV-2**

A. Viral titer in nasal washes of adjuvant-immunized, RBD-nanoparticle IM immunized, or RBD-nanoparticle IM and IN immunized ferrets upon high titer SARS-CoV-2 challenge.

B. Viral titer in lungs of adjuvant-immunized, RBD-nanoparticle IM immunized, or RBD-nanoparticle IM and IN immunized ferrets upon high titer SARS-CoV-2 challenge. Infectious virus titers were measured and shown as mean ± SEM.

